## Supplementary material for "ATR expands embryonic stem cell fate potential in response to replication stress": Ext Figures 1-11

### Extended Data Figure 1

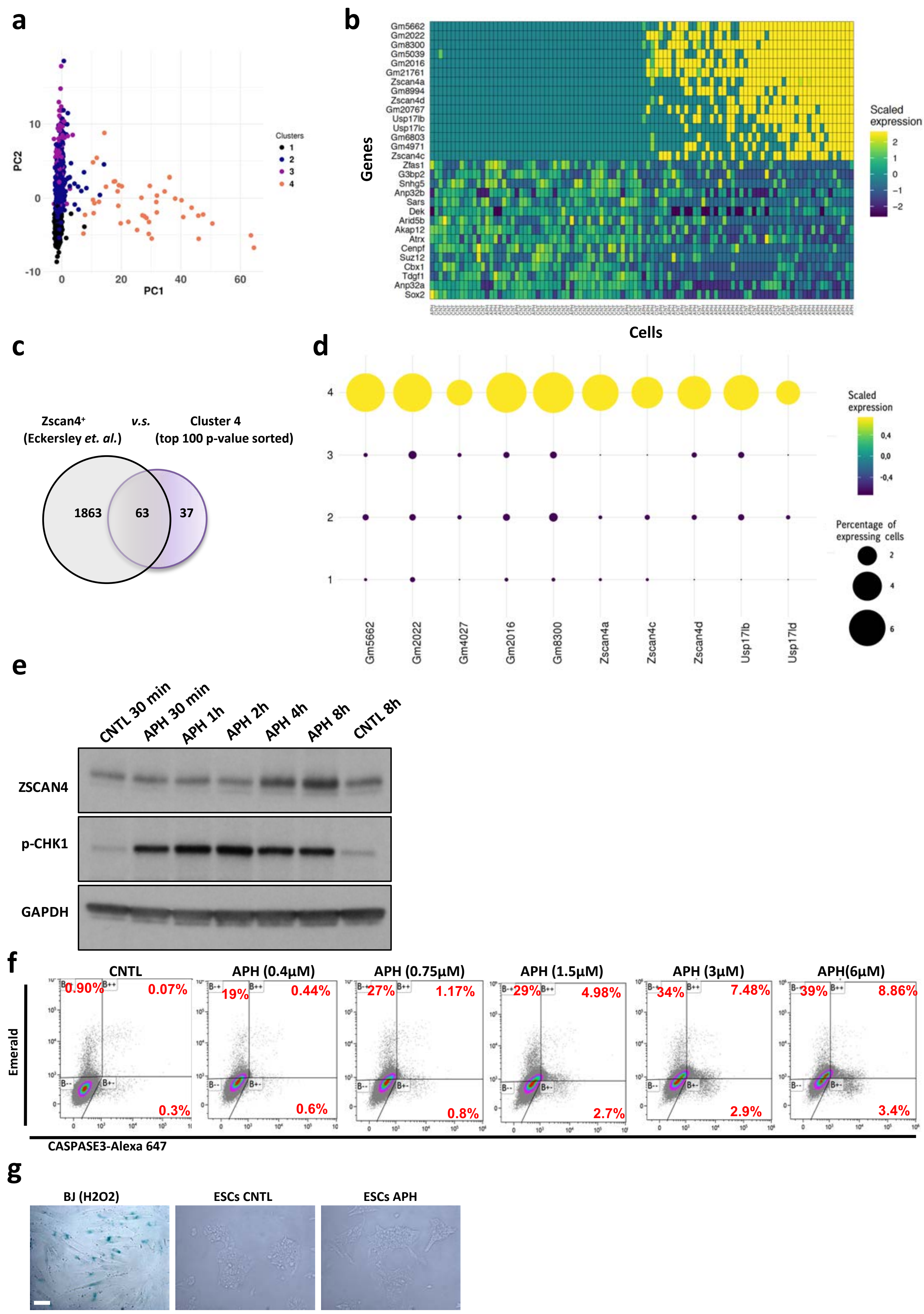

### Extended Data Figure 2

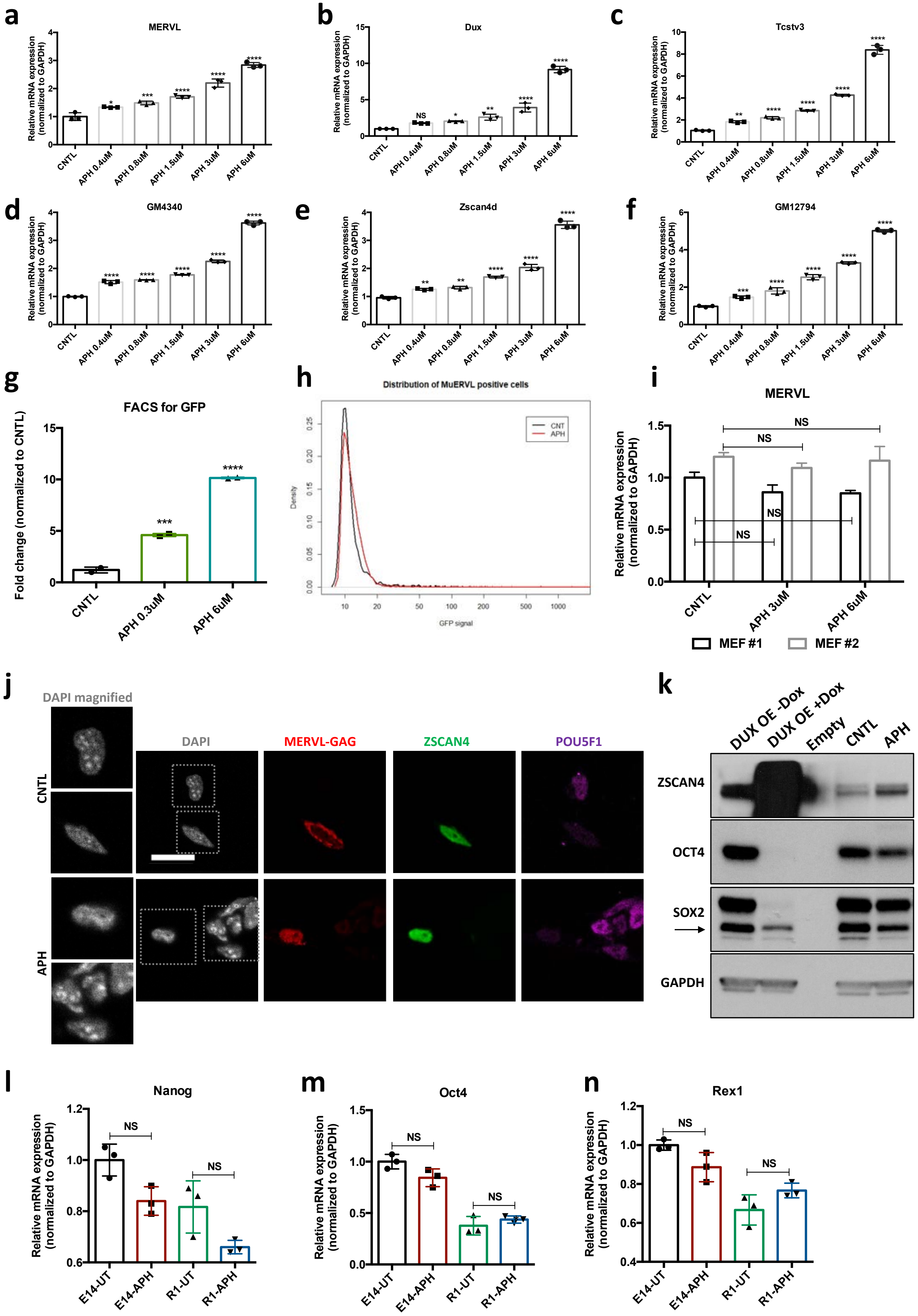

### Extended Data Figure 3

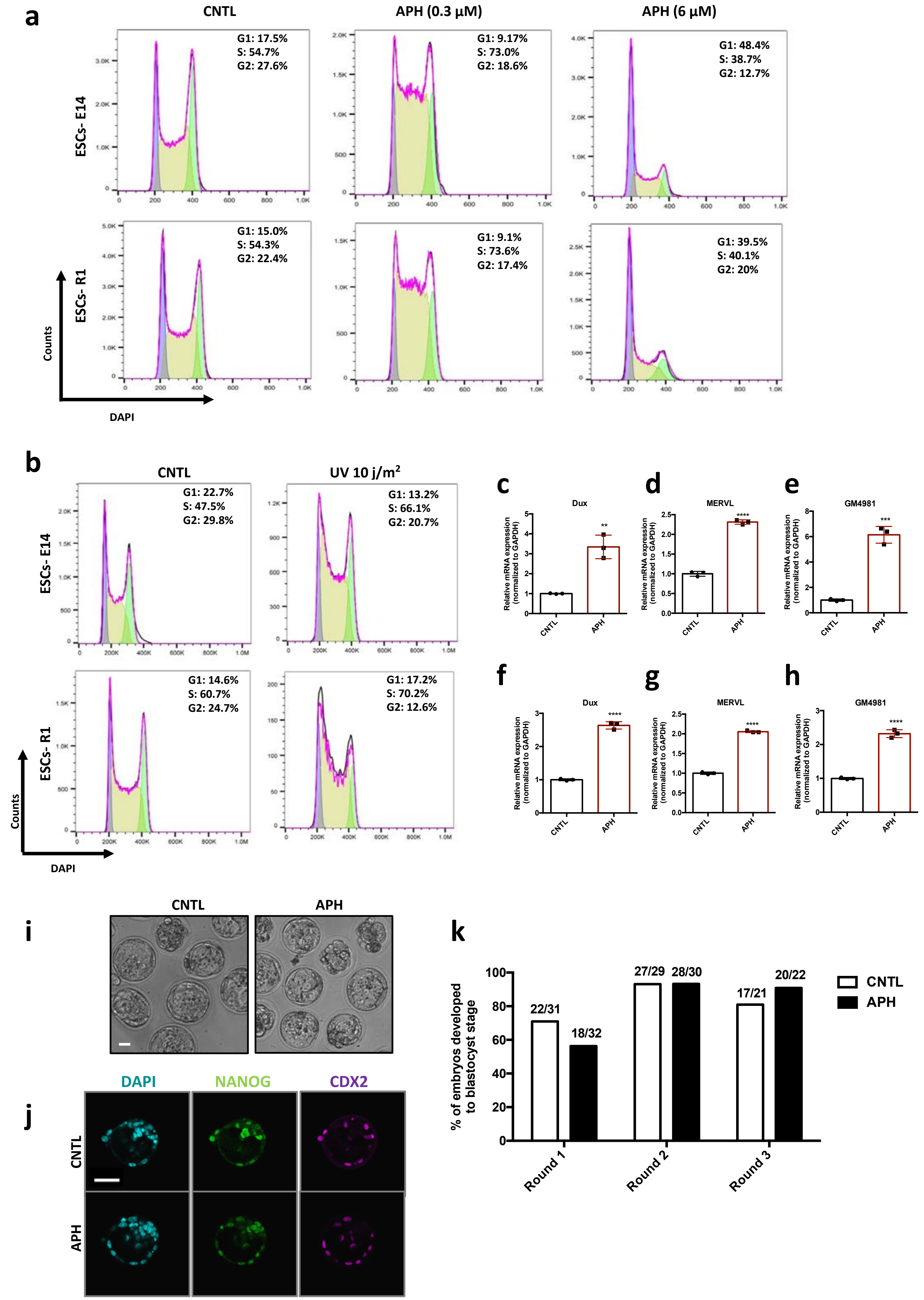

### Extended Data Figure 4

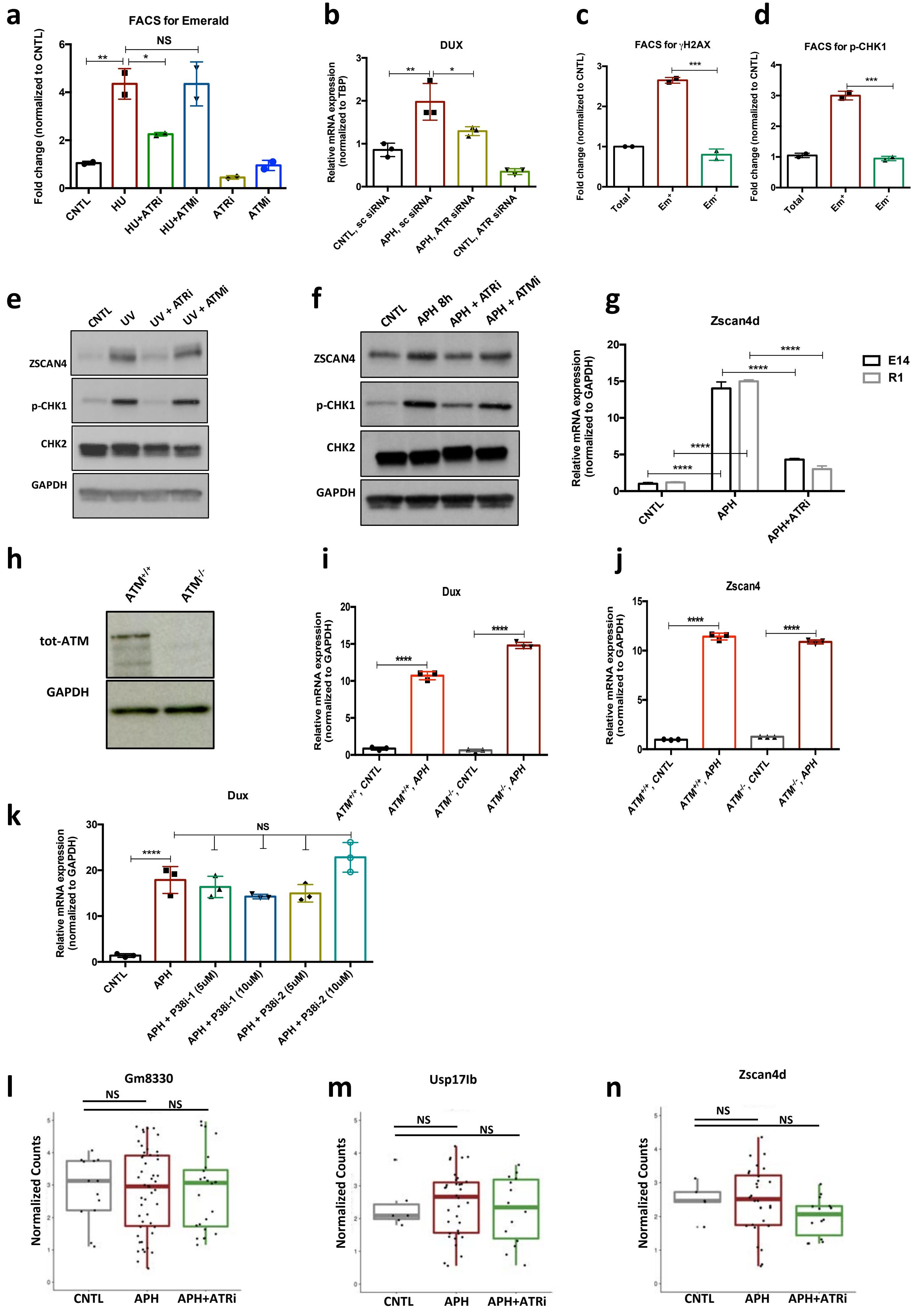

### Extended Data Figure 5

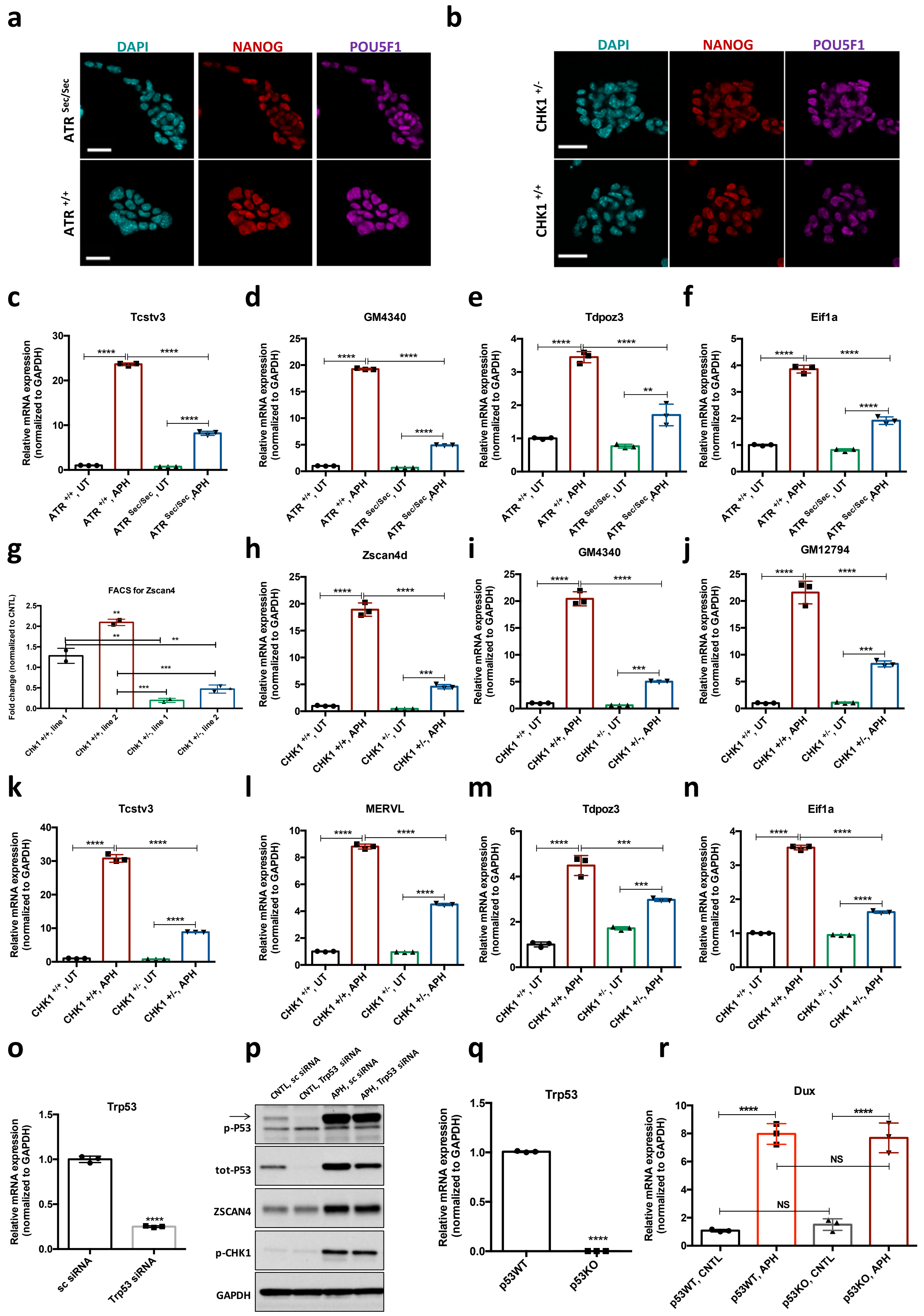

### Extended Data Figure 6

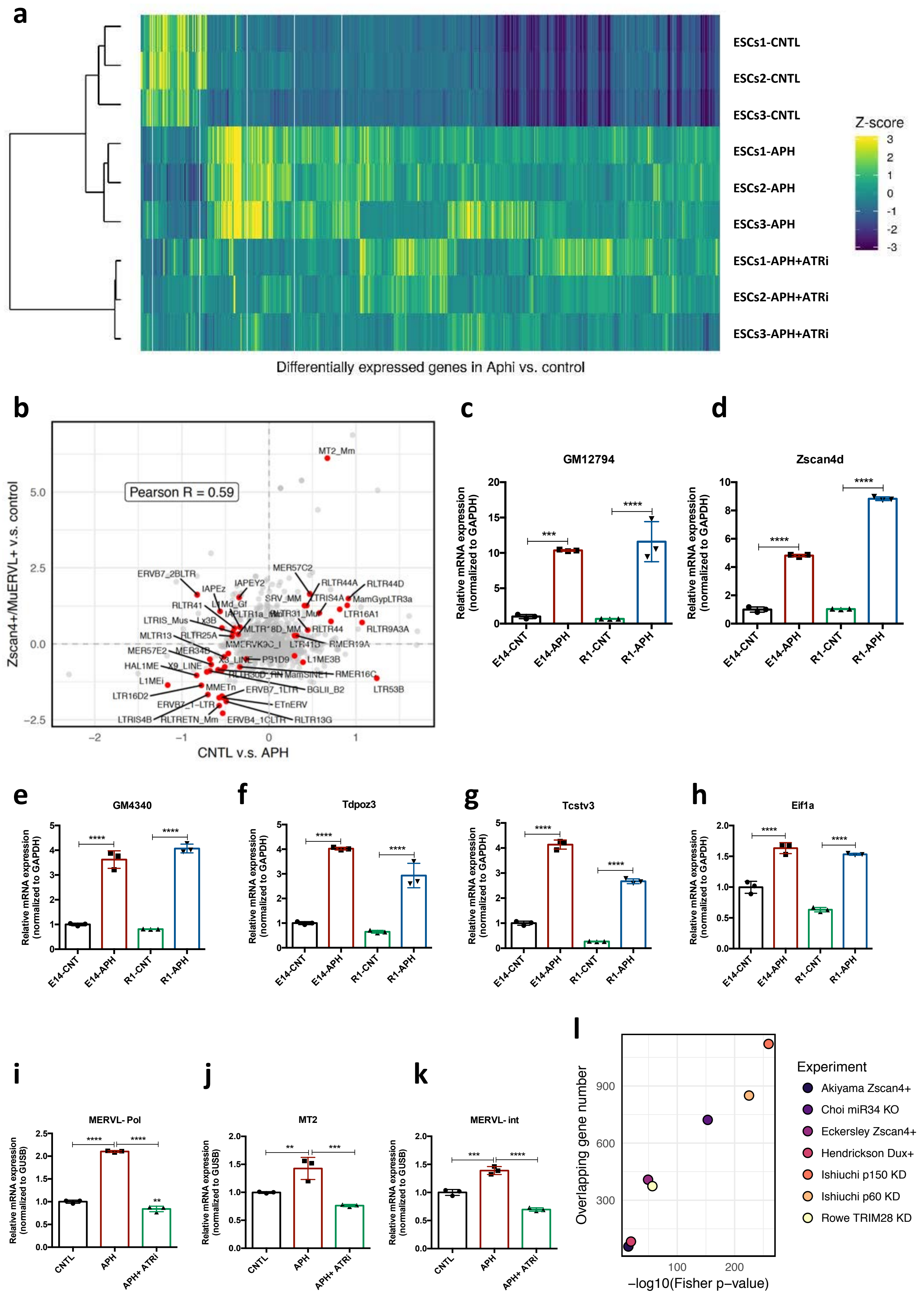

### Extended Data Figure 7

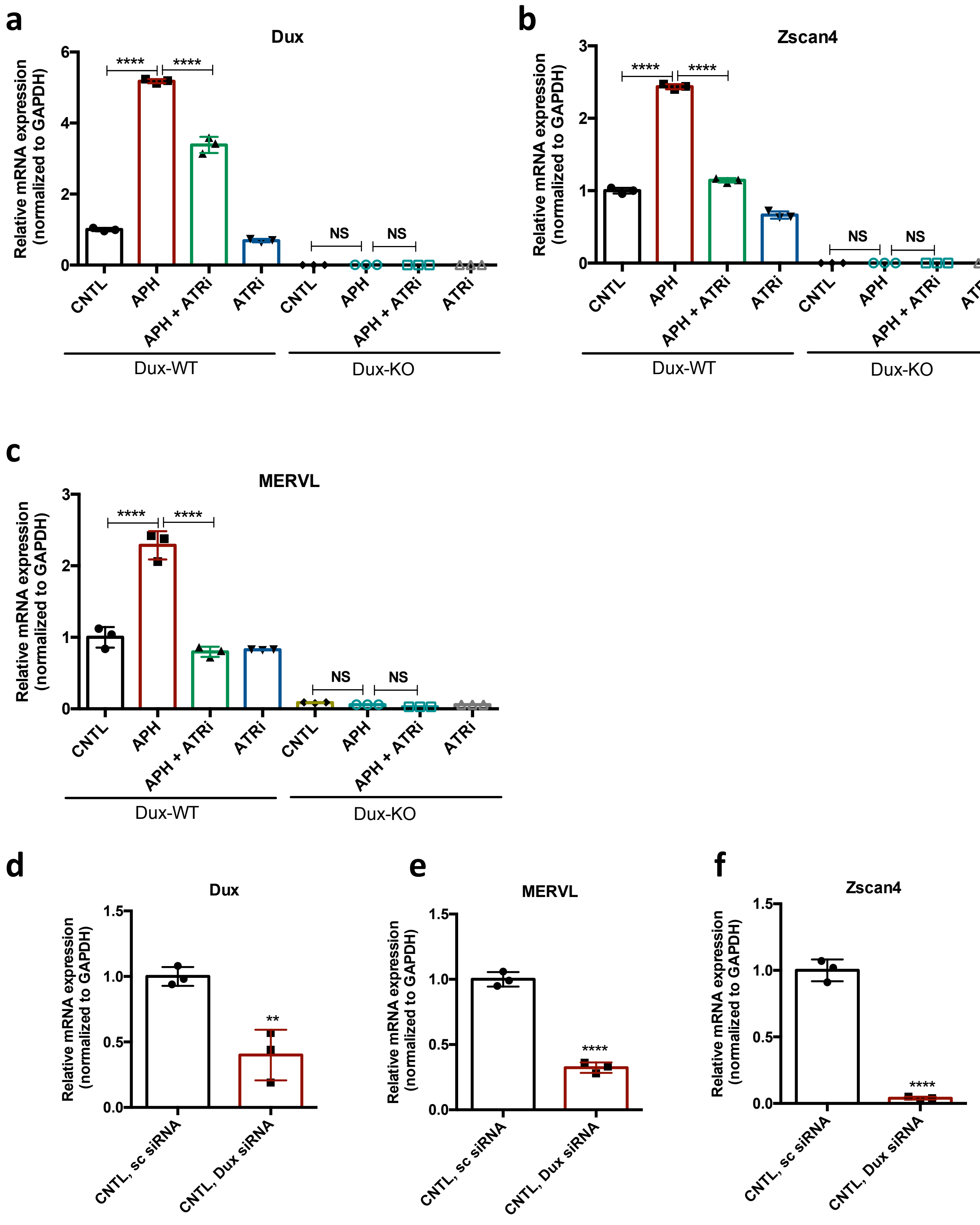

### Extended Data Figure 8

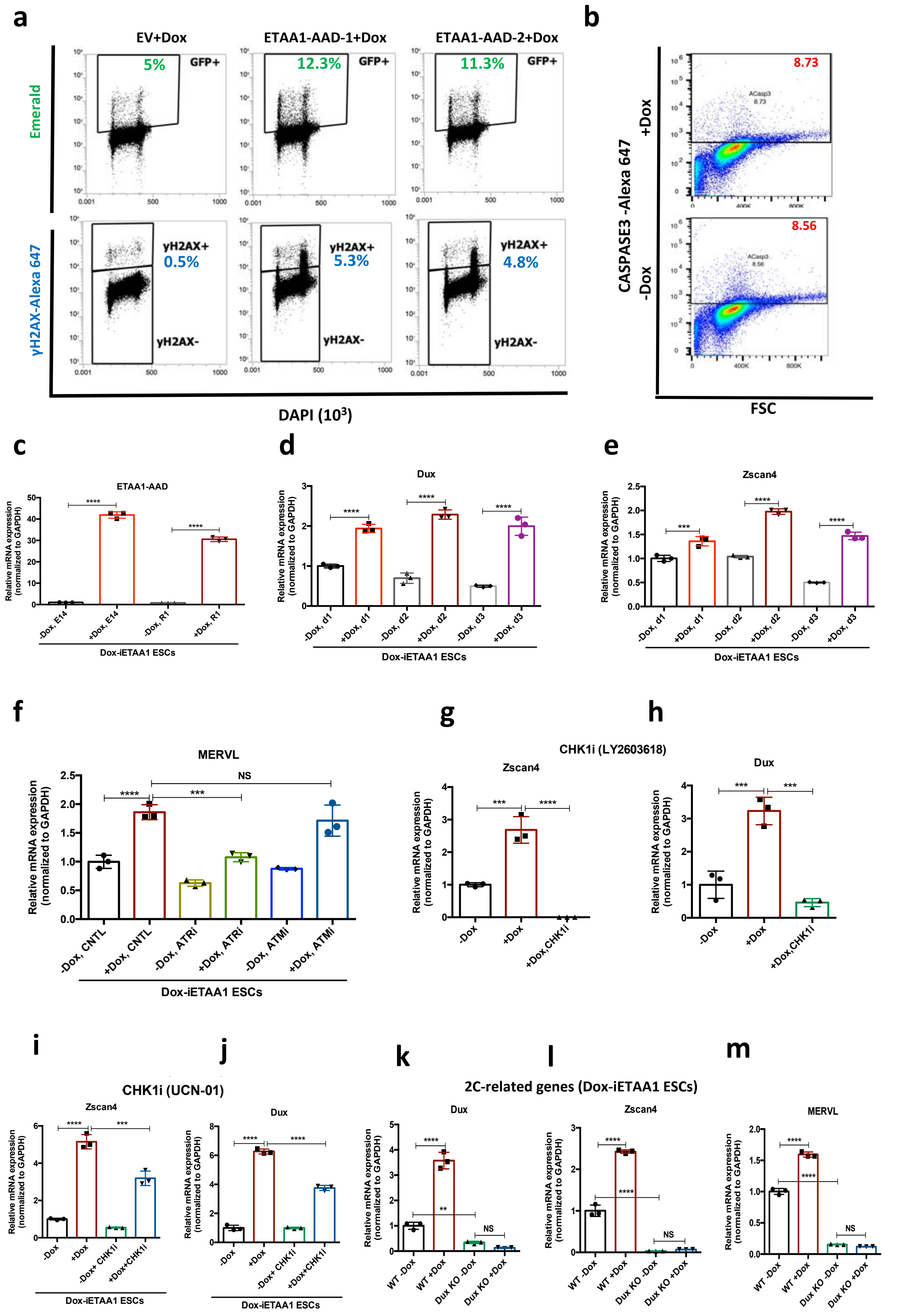

### Extended Data Figure 9

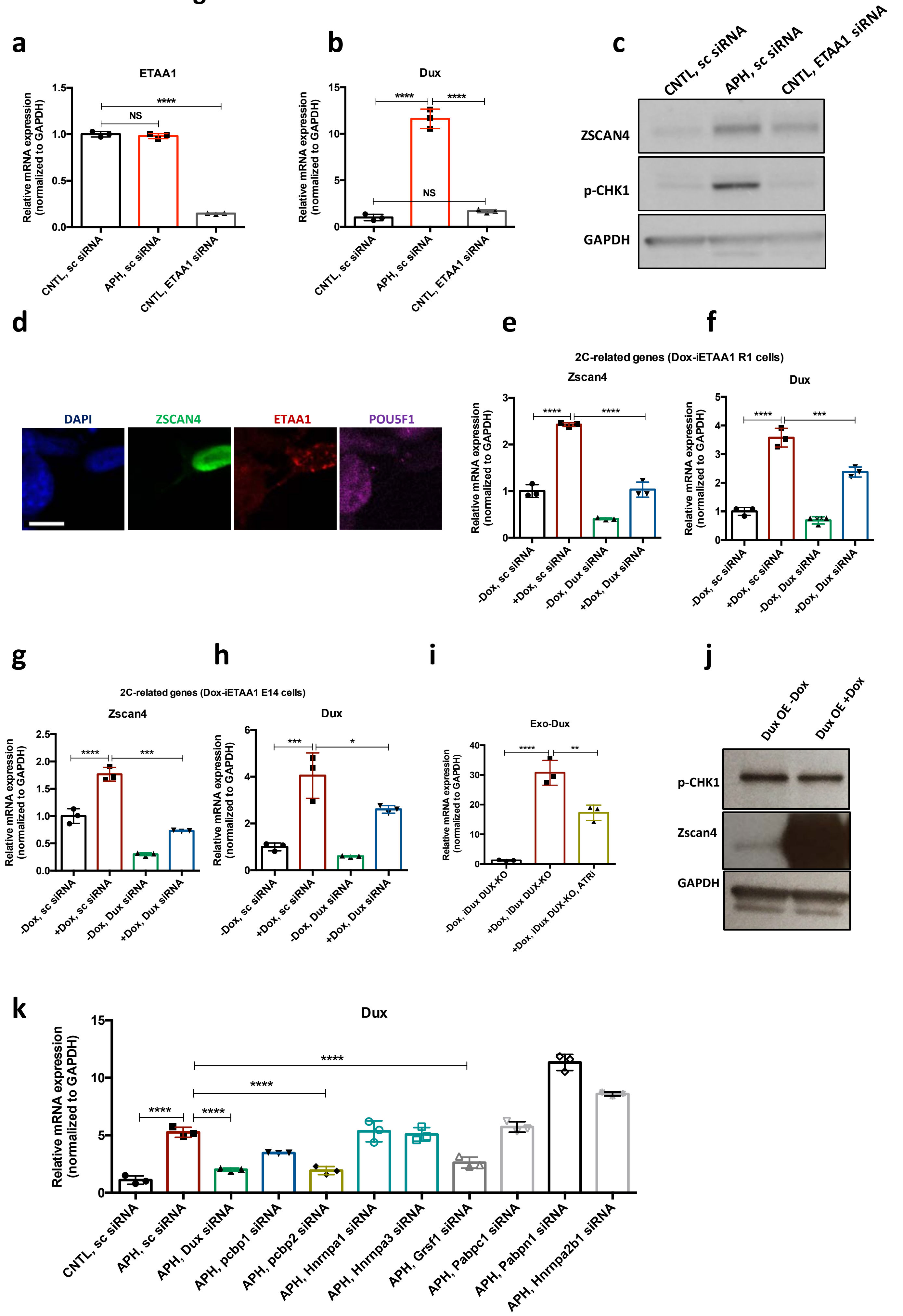

### Extended Data Figure 10

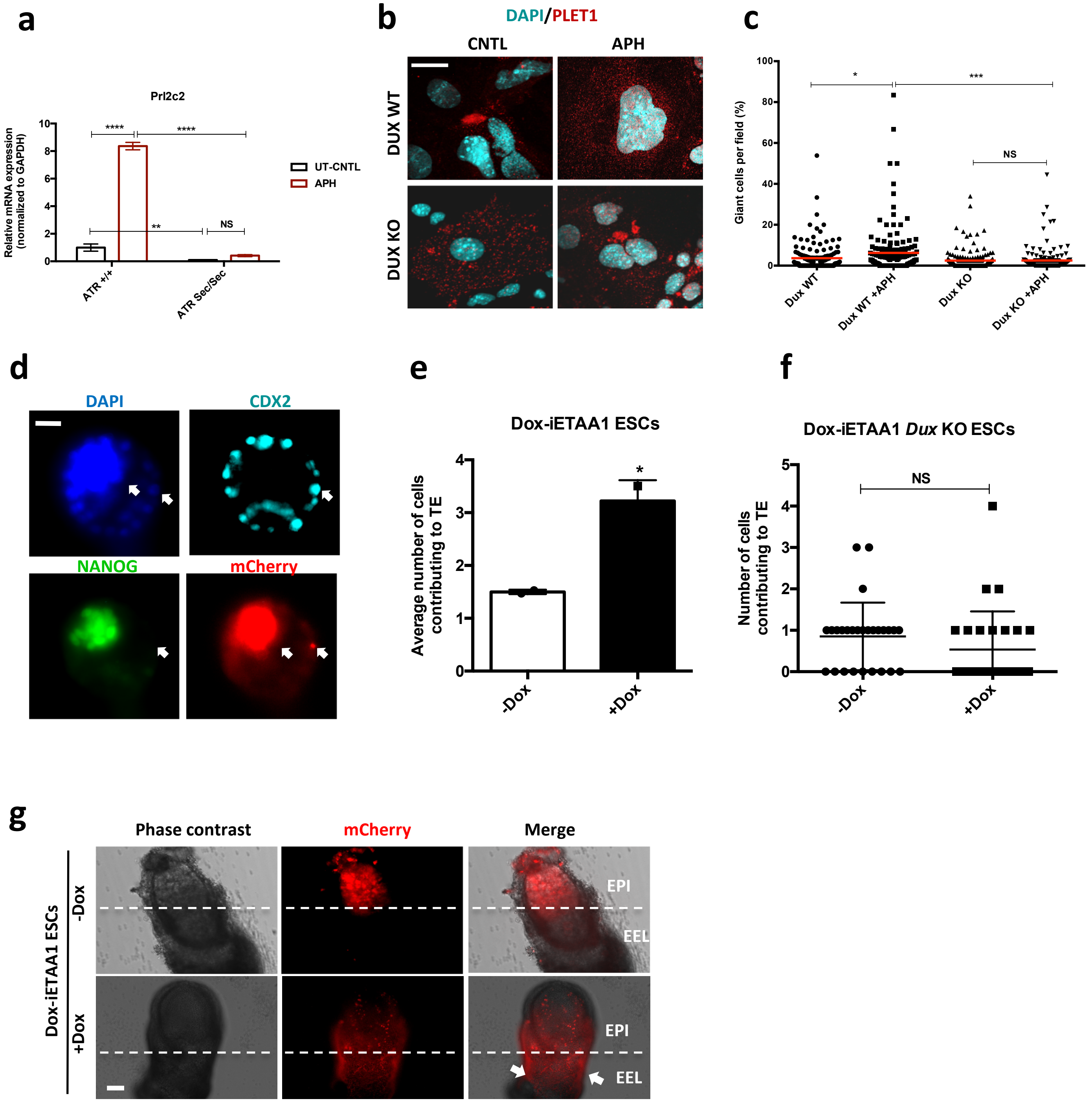

### Extended Data Figure 11

**a**

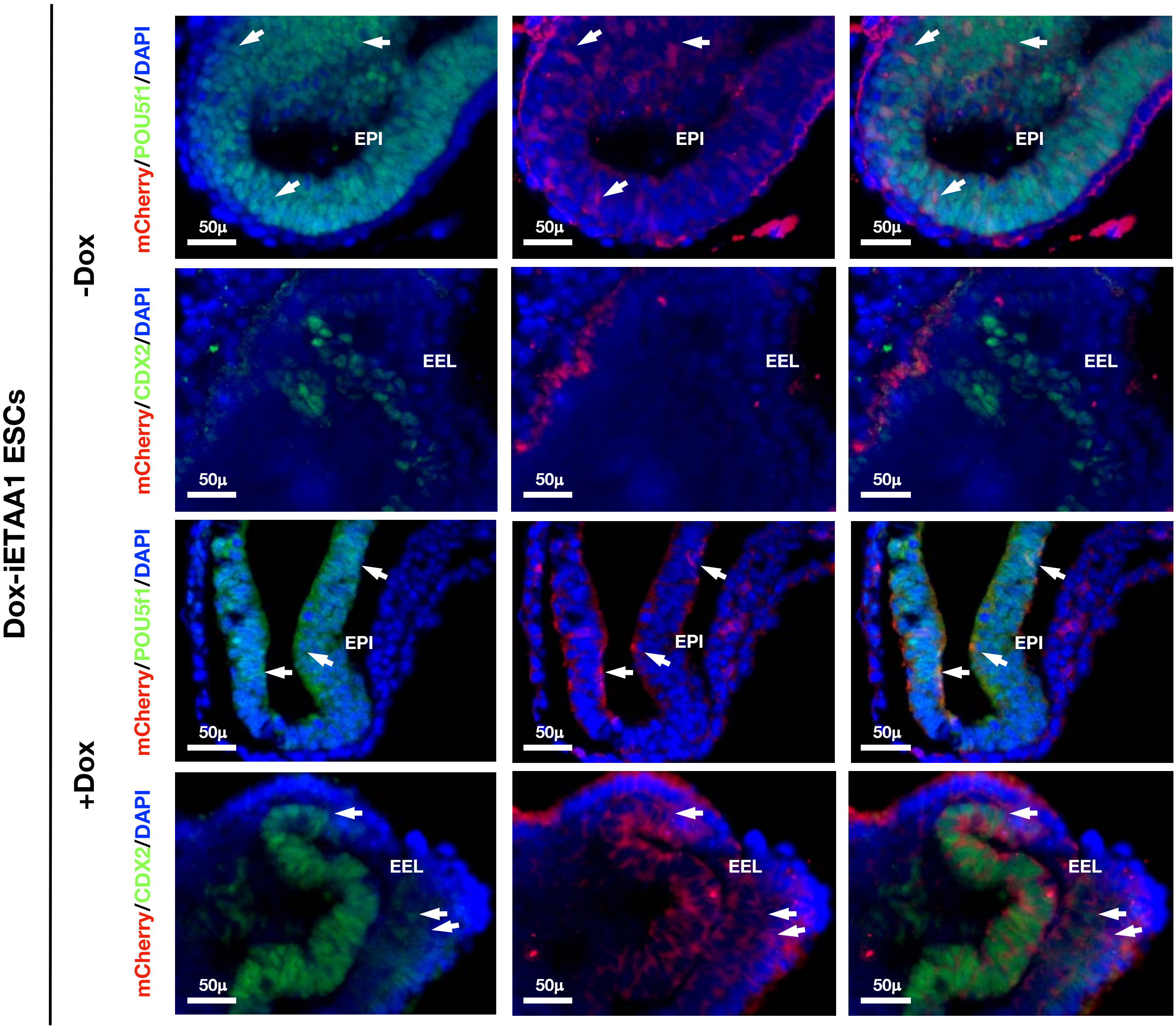


**b**

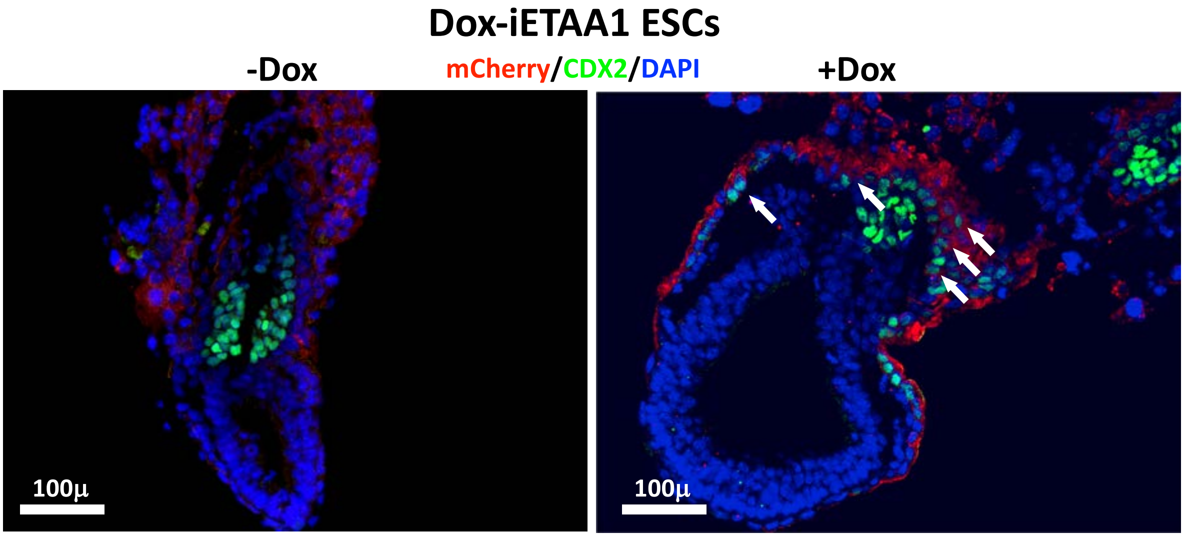
